# Phantasus: web-application for visual and interactive gene expression analysis

**DOI:** 10.1101/2022.12.10.519861

**Authors:** Maksim Kleverov, Daria Zenkova, Vladislav Kamenev, Margarita Sablina, Maxim N. Artyomov, Alexey A. Sergushichev

## Abstract

Transcriptomic profiling became a standard approach to quantify a cell state, which led to accumulation of huge amount of public gene expression datasets. However, both reuse of these datasets or analysis of newly generated ones requires a significant technical expertise. Here we present Phantasus – a user-friendly web-application for interactive gene expression analysis which provide a streamlined access to more than 84000 public gene expression datasets, as well as allows analysis of user-uploaded datasets. Phantasus integrates an intuitive and highly interactive JavaScript-based heatmap interface with an ability to run sophisticated R-based analysis methods. Overall Phantasus allows to go all the way from loading, normalizing and filtering data to doing differential gene expression and downstream analysis. Phantasus can be accessed on-line at https://ctlab.itmo.ru/phantasus or https://artyomovlab.wustl.edu/phantasus or can be installed locally from Bioconductor (https://bioconductor.org/packages/phantasus). Phantasus source code is available at https://github.com/ctlab/phantasus under MIT licence.

## 1 Introduction

Transcriptomic profiling is an ubiquitous method for whole-genome level profiling of biological samples [Stark et al., 2019]. Moreover, deposition of these data into one of the public repositories became a standard in the field, which led to accumulation of huge amount of publicly available data. The most significant example is NCBI Gene Expresision Omnibus (GEO) project [Barrett et al., 2012], which stores information of more than 180000 studies.

Sharing of transcriptomic data opens up possibilities for reusing them: instead of carrying out a costly experiment, a publicly available dataset can be used, thus decreasing the cost and accelerating the research [Byrd et al., 2020]. However, the standard approach for gene expression analysis requires a significant technical expertise. In particular, many analysis methods are implemented in R as a part of Bioconductor project ecosystem [Gentleman et al., 2004], and thus one has to have programming skills in R to use them. On the other hand, domain knowledge is beneficial to improve quality control of the data, which is especially important when working with the publicly available data, as well as generation of biological hypotheses [Wang et al., 2016].

A number of applications have been developed with the aim to simplify analysis of transcriptomic datasets (see Supplementary File 1 for details). In particular, web-based applications remove the burden of set-up and configuration from the end users, thus lowering the entry threshold. Shiny framework [Chang et al., 2022] revolutionized the field as it became easy to create a web interface for R based pipelines, which led to a significant growth of web-applications for gene expression analysis [Ge et al., 2018, Iacoangeli et al., 2022, Mahi et al., 2019, Nelson et al., 2016]. However, such applications generally have limited interactivity due to mainly server-side computations. Shiny-independent applications can be more interactive, but they suffer from lack of native R support and require reimplementation of existing methods from scratch [Gould, 2016, Alonso et al., 2015].

Here we present Phantasus: a web-application for gene expression analysis that integrates highly interactive client-side JavaScript heatmap interface with an R-based backend. Phantasus allows to carry out all major steps of gene expression analysis pipeline: data loading, annotation, normalization, clustering, differential gene expression and pathway analysis. Notably, Phantasus provides a streamlined access to more than 84000 microarray and RNA-seq datasets from Gene Expression Omnibus database, simplifying their reanalysis. Phantasus can be accessed on-line at https://ctlab.itmo.ru/phantasus or at https://artyomovlab.wustl.edu/phantasus or can be installed locally from Bioconductor. Phantasus is open source and its code is available at https://github.com/ctlab/phantasus under MIT licence.

## 2 Results

### 2.1 Phantasus web-application

We developed a web application called Phantasus for interactive gene expression analysis. Phantasus integrates JavaScript-rich heatmap based user interface originated from Morpheus [Gould, 2016] with an R back-end via OpenCPU framework [Ooms, 2014]. Heatmap graphical interface provides an intuitive way to manipulate the data and metadata: directly in a web-browser the user can create or modify annotations, edit color schemes, filter rows and columns, and so on. On the other hand the R back-end provides a way to easily run a multitude of computational analysis methods available as R packages. All together this architecture (Figure 1) provides a smooth experience for doing all common analysis steps: loading datasets, normalization, exploration, visualization, differential expression and gene set enrichment analyses.

**Figure 1:**
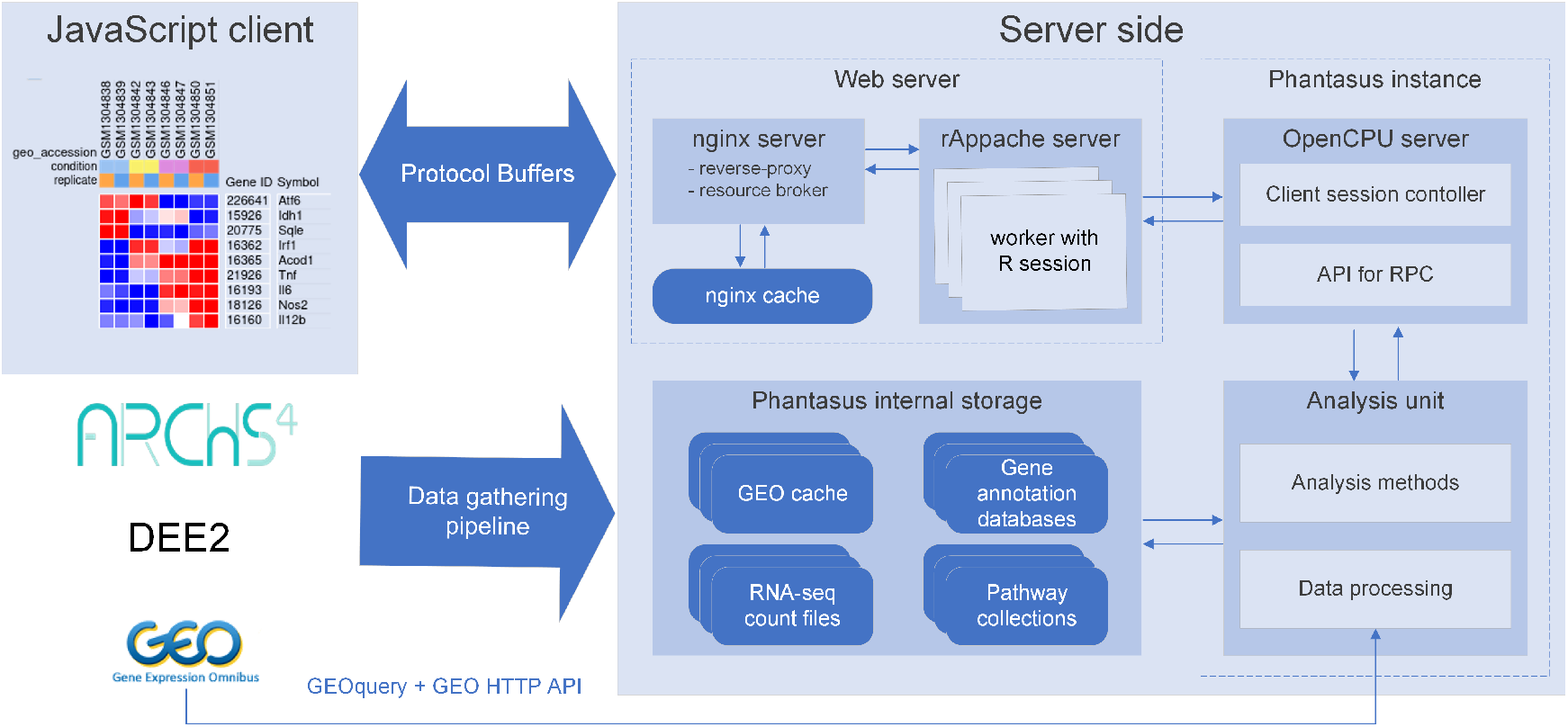
Overview of Phantasus architecture. The front-end interface is a JavaScript application, that requests the web-server to load the data and perform resource-consuming tasks. The core element of the back-end is the OpenCPU-based server which triggers execution of R-based analysis methods. Protocol Buffers are used for efficient client-server dataset synchronization.

Several options to load the gene expression data into Phantasus are available. First, datasets from Gene Expression Omnibus [Barrett et al., 2012] can be loaded by their identifier. Phantasus supports microarray datasets, which are loaded directly from GEO, as well as RNA-sequencing datasets, for which counts data from third-pary databases are used (see section Section 2.3 for details). Second, datasets can be loaded from a gene expression table file in GCT, TSV and XLSX formats. Finally, a set of curated datasets are available directly from the home page.

A number of methods can be used to prepare, normalize and explore the gene expression table. In particular, it is possible to aggregate microarray probe-level data to gene levels, transform and filter the data, do a principal component analysis (PCA), do a k-means or hirarchical clustering, etc. These tools allows to do a thorough quality control of the dataset and remove the outliers if they are present.

When the dataset is properly filtered and normalized, differential expression analysis using limma [Ritchie et al., 2015] or DESeq2 pipelines [Love et al., 2014] can be carried out. These results can then be used with other web-services for downstream analysis, with shortcuts for pathway analysis with Enrichr [Kuleshov et al., 2016] and metabolic network analysis with Shiny GAM [Sergushichev et al., 2016]. Additionally, gene set enrichment analysis can be done directly in Phantasus as implemented in fgsea package [Korotkevich et al., 2021].

All of the plots produced by Phantasus during the data exploration and analysis can be exported as vector images in SVG format. This includes heatmaps, PCA plots, gene profiles, enrichment plots etc. The obtained images can be used for publications as is or adjusted in a vector graphics editor.

Another option for presenting final or intermediate results is a session link sharing. When a link is generated, a snapshot with the current dataset and its representation: annotations, color scheme, sample dendrograms, etc, is saved on the server. The link can be shared with other users, and, when opened, restores the session.

### 2.2 Stand-alone phantasus distribution

Aside from using the two official mirrors: https://ctlab.itmo.ru/phantasus and https://artyomovlab.wustl.edu/phantasus, there is a possibility to set up phantasus locally. Phantasus can be installed as an R package from Bioconductor (https://bioconductor.org/packages/phantasus) or loaded as a Docker image (https://hub.docker.com/r/asergushichev/phantasus). In both cases almost all of the Phantasus functions will be available from the start.

Some of Phantasus features require additional server-side set up. Extended support of GEO datasets requires preprocessed expression matrices and platfrom annotations. Identifier mapping requires organism annotation databases. Pathways enrichment requires pathway databases. For the initial set up, all these files can be downloaded from https://ctlab.itmo.ru/files/software/phantasus/minimal-cache.

Important feature of a stand-alone version of Phantasus is an ability to share manually curated datasets. Similar to Phantasus session link sharing, one can generate a named session consisting of a dataset and its visual representation. Link to this named session (e.g. https://ctlab.itmo.ru/phantasus/?preloaded=GSE53986.Ctrl.vs.LPS) can then be shared for the other users to view. Such predictable display of the data can be pariticularly useful in a publication context.

### 2.3 Available datasets

Phantasus provides a streamlined access to more than 84379 GEO datasets. For these datasets the expression values and gene identifiers (Entrez, ENSEMBL or Gene Symbol) are readily available (Figure 2). Moreover, these datasets are used to populate initial Phantasus cache, and thus they have low loading times.

**Figure 2:**
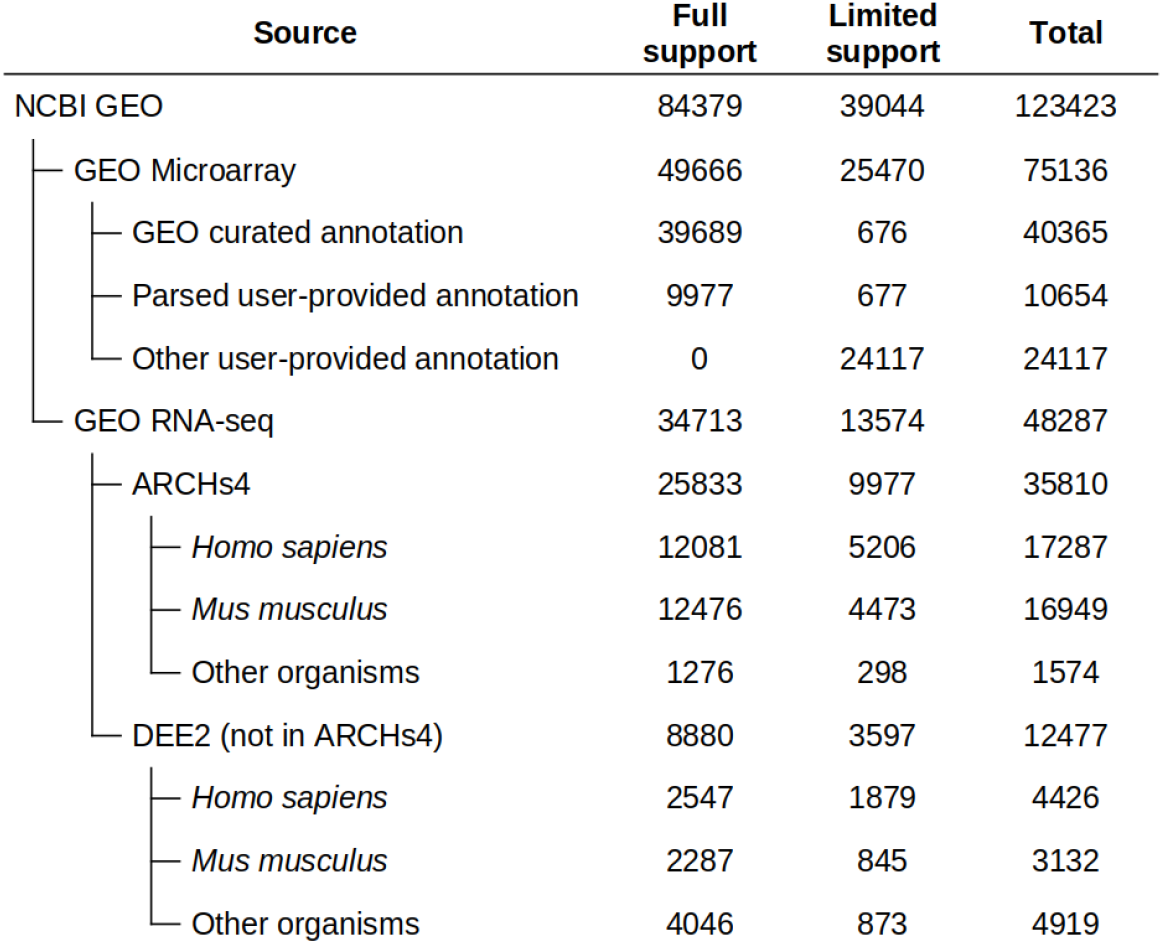
Dataset availablility in Phantasus. For fully supported datasets gene expression data is accompanied by gene annotations in a standardized format. Limited support datasets have either incomplete gene expression matrix or gene annotations.

**Figure 3:**
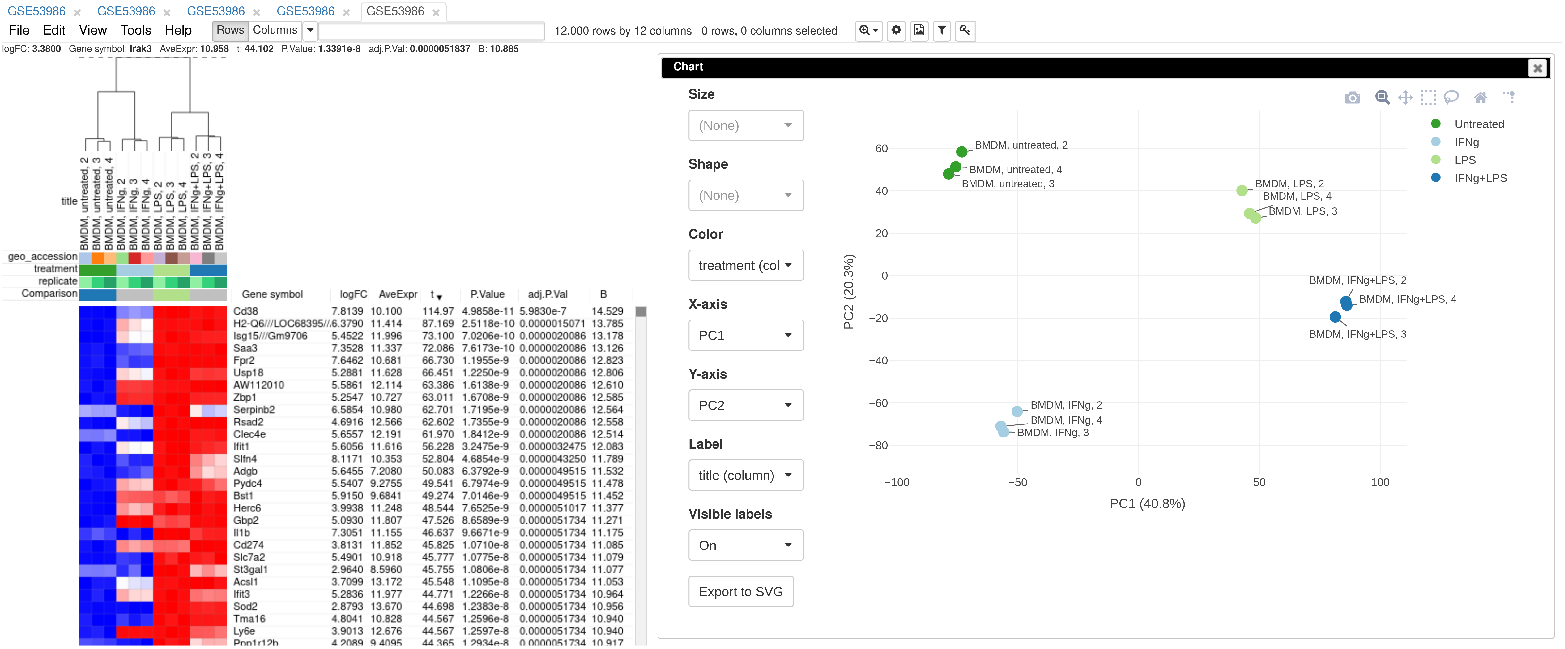
Example of analysed dataset GSE53986 with normalized gene expression values, filtered outliers, hierarchically clustered columns, and rows annotated with differential expression analysis between untreated and LPS-treated macrophages

From these 8484379 datasets 49666 are microarrays based on 2767 platforms. For 1347 platforms GEO databses have a machine-readable annotations in the *annot.gz* format with Entrez gene and Gene symbol columns, which correspond to 39689 datasets. The remaining 9977 datasets are obtained from platforms that do not have a GEO-provided annotation. For these 1420 platforms we have automatically marked up user-provided annotations in *SOFT* format to extract gene identifiers and convert the annotations into *annot.gz* format.

RNA-seq subset of the datasets with a streamlined access consits of 34713 datatasets. As GEO does not store expression values for RNA-seq datasets, we rely on other databases for the expression data. The first-priority database for RNA-seq gene counts is ARCHS4 (Human, Mouse and Zoo versions), which covers 25833 datasets. The other source is DEE2 database (human, mouse and other available organisms), which covers an additional 8880 datasets. DEE2 database contains transcript-level quantification, so it has been preprocessed to sum read counts into gene-level tables.

## 3 Implementation

### 3.1 Web application architecture

Phantasus is a web-application that combines interactive graphical user interface with an access to a variety of R-based analysis methods (Figure 1). The front-end is JavaScript-based, and is derived from Morpheus web-application for matrix visualization and analysis [Gould, 2016]. The back-end is written in R, with an OpenCPU server [Ooms, 2014] translating HTTP-queries from the client into R procedure calls.

The JavaScript client is responsible for the matrix visualization, as well as certain analysis methods. In particular, steps like subsetting the dataset, working with annotations, basic matrix modification (e.g., log-transformation, scaling, etc) have client-side implementation. Furthermore, the client supports additional visualization methods such as row profile plots, volcano plot, and others.

The analysis methods that require external data or algorithms are implented in the form of the `phantasus` R package to be carried out on the server side. The operations include differential gene expression analysis, principal component analysis, pathway analysis, and others. Commonly, these methods rely on functions which are already available in the existing R packages, for such methods only wrapper R functions are implemented.

OpenCPU server is a core component of the Phantasus back-end. The server provides an HTTP API for calling computational methods implemented in R. For each call OpenCPU creates a new R environment with the required data, in which the method is then executed. OpenCPU can manage these R environments both in a standard single-user R session, and, with the help of rApache, in a multi-user manner inside an Apache web-server.

The transfer of large objects between the server and the client exploits a binary Protobuf protocol. The Phantasus back-end uses protolite R package [Ooms, 2021] for object serialization and deserialization. The front-end relies on protobuf.js module [Coe, 2020].

For the further performance improvement Nginx server is used to wrap OpenCPU server. Nginx server caches the results of the OpenCPU method calls. If the same method with the same data is called again the cached result can be returned without any additional computations. Furthermore Nginx is used to serve static content and to manage permissions.

### 3.2 Data sources and data gathering

The main data source for Phantasus is NCBI GEO database [Barrett et al., 2012]. All of the GEO datasets are identified by a GSEnnnnn accession number (with a subset of the datasets having an additional GDSnnnnn identifier). However, depending on the type of the dataset the processing procedure is different.

The majority of gene expression datasets in GEO database can be divided into two groups: microarray data and RNA-seq data. While the experiment metadata is available for all of the datasets, the expression matrices are provided only for the microarray datasets. Phantasus relies on GEOQuery package [Davis and Meltzer, 2007] to load the experiment metadata (for all of the datasets) and expression matrics (for microarray datasets) from GEO.

When a GEO RNA-seq dataset is requested by the user, Phantasus refers to precomputed gene counts databases availble in the internal storage. In particular, data from ARCHS4 [Lachmann et al., 2017] and DEE2 [Ziemann et al., 2019] projects are used. Both of these projects contain gene counts and metadata for RNA-seq samples related to different model organisms including but not limited to mouse and human. For any requested RNA-seq dataset the gene counts are loaded from a single database, whichever covers the highest number of samples.

Next, Phantasus stores gene annotation databases which are used to map genes between different identifier types. Theese databases are stored in sqlite format compatible with *AnnotationDbi* R package [Pagès et al., 2022]. Currently only human and mouse databases are available, which are based on *org.Hs.eg.db* and *org.Mm.eg.db* R packages respectively.

Pathway databaes are stored to be used for gene set enrichment analysis. Currently gene set collections include GO biological processes database [Ashburner et al., 2000], Reactome database [Gillespie et al., 2021] and MSigDB Hallmark database [Liberzon et al., 2011] for human and mouse.

Finally, for a faster access, Phantasus dataset cache is automatically populated by a large compendium of datasets. The automatic piepline is Snakemake-based and consists of four steps. First of all the pipeline converts DEE2 files to ARCHs4-like HDF5 files. During this procedure transcript expression provided by DEE2 is summed up to gene level. Second step checks for which microarray platforms GEO contains a curated machine-readable annotation in *annot.gz* format. Third step tries to generate the machine-readable annotation for the rest of microarray platforms from the annotations available in the *SOFT* format. Currently, this step procduces an additional 1300 *annot.gz* files. The last step goes over all of the microarray datasets with a machine-readable annotation and over all of the RNA-seq datasets with the counts available in ARCHS4 or DEE2. For each such dataset the cached entry with all of the data and metadata is created and stored.

A snapshot of Phantasus internal storage is available at https://ctlab.itmo.ru/files/software/phantasus/minimal-cache. It contains preprocessed count files, automatically marked-up annotations, gene and apthways databases. This snapshot can be used for a local Phantasus set up.

## 4 Case study

### 4.1 Basic usage

To illustrate the basic usage of Phantasus we will consider dataset GSE53986 [Noubade et al., 2014] from GEO database. This dataset conists of 16 samples of bone marrow derived macrophages, untreated and treated with three stimuli: LPS, IFNg and combined LPS+INFg. The gene expression was measured with Affymetric Mouse Genome 430 2.0 Array array. Here we give an overview of the steps, the full walk-through for the analysis is available in Supplementary File 2.

As the first step of the analysis the dataset can be loaded and normalized. The dataset is loaded straightforwardly by the *GSE53986* identifier. Because this is a microarray dataset, internally the gene expression values are obtained from GEO. In this particular case the expression values have not been normalized, but it can be done in Phantasus. From the available normalization options we select log2 scaling and quantile normalization. Further, we can aggregate microarray probe-level expression values into gene-level expression. We chose *Maximum Median Probe* method which retains only a single probe per gene, the one that has highest median expression value. Finally, we can filter out lowly expressed genes, for example, by keeping only the top 12000 expressed genes.

After the normalization step we can apply a number of exploratory techiniques. In particular we can do a Principal Component Analysis, k-means gene clustering and hierarchical clustering of the samples. From these analysis we can discover, that there is an overall good concordance between the replicates of the same treatment, with an exception of the first replicate in each group. We can conclude that these samples are outliers and remove them before the downstream analysis.

Finally, we can do a comparison between the sample groups, for example by comparing untreated and LPS-treated samples. As the data has been normalized, we can apply *limma* for differential gene expression analysis. The result appears as additional gene annotation columns: P-values, log-fold-changes and other statistics. Next we can use differential expression results for a pathway enrichment analysis: for example, we can use R-based gene set enrichment analysis vai *fgsea* package or we can use external tools, such as Enirhcr.

### 4.2 Data reanalysis

To highlight Phantasus ability to reanalyze publicly available data in a context of a biological study let us consider a study by Mowel and colleagues [Mowel et al., 2017]. The study considers a genomic locus *Rroid* linked by the authors to homeostasis and function of group 1 innate lymphoid cells (ILC1). The authors hypothesized that *Rroid* locus controls ILC1s by promoting the expression of Id2 gene, a known regulator of ILCs. To confirm this hypothesis the authors generated an Id2-dependent gene signature based on an existing transcriptomic data [Shih et al., 2016] and showed its deregulation in *Rroid* deficient cells.

The descibed above computational analysis linking Rroid and Id2 can be replicated in Phantasus in a straightforward way (see Supplementary File 3 for the detailed walk-through).

First, we can open GEO dataset GSE76466 [Shih et al., 2016], containing gene expression data for Id2 deficient NK cells. Notably, GSE76466 is an RNA sequencing dataset, whithout gene expression values stored directly in GEO database, however Phantasus loads the dataset leveraging precomputed expression values from ARCHS4 project [Lachmann et al., 2017]. Then we can run differential gene expression analysis with DESeq2, comparing Id2-deficient and wild-type NK cells. Id2-dependent gene signature can be obtained by sorting the genes by *stat* column.

Second, RNA-sequencing dataset GSE101459 [Mowel et al., 2017], generated by Mowel and col-leagues for Rroid-deficient NK cells, can also be opened in Phantasus. There we can do differential gene expression analysis with DESeq2 and remove lowly expressed genes. Finally we can enter the generated Id2-dependent gene signature into Phantasus gene search field and use GSEA plot tool to obtain an enrichment plot, similar to one presented by Mowel and colleagues, confirming a potential regulation via Id2.

## 5 Conclusion

Phantasus is a tool for visual and interactive gene expression analysis that allows in an easy and streamlined manner to go from loading, normalizing and filtering data to differential gene expression and downstream analysis. Additionally, due to its tight integration with R environment, Phantasus can be extended with other analysis methods, in particular the ones available at Bioconductor. Phantasus can be both used on-line at https://ctlab.itmo.ru/phantasus or be installed locally from Bioconductor.

## 6 Acknowledgements

The project was supported by Ministry of Science and Higher Education of the Russian Federation (Priority 2030 Federal Academic Leadership Program).

