## Supplementary File 1 for "Phantasus: web-application for visual and interactive gene expression analysis"

**Supplementary File 1.** Comparison of gene expression analysis platforms

In this section we consider existing software platforms for gene expression analysis. While they share common principles, the details of implementation vary, which affects their usability. For the comparison we considered three aspects: 1) support for gene expression analysis steps, 2) data availability, and 3) user experience.

The general gene expression analysis workflow can be roughly divided into three stages: data preprocessing, exploratory analysis and differential gene expression with pathway analysis. The Table 1 contains an overview of support of the key analyis steps implemented in the considered tools. At the data preprocessing stage many tools allow to load unnormalized gene expression matrices and provide a variety of matrix normalization approaches such as log-transformation, quantile normalization, scaling and others. After the dataset is normilized, some of these tools provide instruments for dataset exploraion like principal component analysis and row/column clustering. Gene expression analysis usually ends with determining differentially expressed genes and pathway analysis to explain observed difference. While the differential expression is available almost everywhere the pathway analysis, can be caried out only by about a half of the tools. In this half the most notable tools are Gene Pattern, Babelomics, iDep, EXPANDER and GEOexplorer which support all the considered steps.

Table 1: Gene expression analysis key steps

| Application | Normalization | PCA | Clustering | Differential expression | Pathway analysis |
|---|---|---|---|---|---|
| Gene pattern [2006] | + | + | + | + | + |
| GEO2R [2012] | - | - | - | + | - |
| BicOverlapper2 [2014] | - | - | - | + | + |
| Babelomics [2015] | + | + | + | + | + |
| Morpheus [2016] | + | - | + | - | - |
| START [2016] | + | + | - | + | - |

| Application | Normalization | PCA | Clustering | Differential expression | Pathway analysis |
|---|---|---|---|---|---|
| BioJupies [2018] | + | + | + | + | + |
| iDEP [2018] | + | + | + | + | + |
| Degust [2019] | - | + | - | + | - |
| GREIN [2019] | + | + | - | + | - |
| EXPANDER [2019] | + | + | + | + | + |
| RaNa-seq [2019] | - | + | - | + | + |
| RNAdetector [2021] | - | - | - | + | - |
| GEOexplorer [2022] | + | + | + | + | + |
| **Phantasus** | + | + | + | + | + |

The huge difference between tools appears in the supported data sources and the availability of datasets (Table 2). In addition to a simple loading of gene expression matrices in tabular formats, which is a comon pratice, few tools allow to load publicly available datasets from external sources, in particular from Gene Expression Omnibus (GEO) database. However, depending on the type of the GEO dataset: microarray or RNA-seq, their support varies among the considered tools. GEO database does not store gene expression matrices for RNA-seq datasets, thus tools like GREIN and RaNa-seq incorporate allignment and quatification of the raw data into their pipeline. Alternatively, precomputed count matrices from other projectes, such as ARHCs4 [Lachmann et al., 2017] and DEE2 [Ziemann et al., 2019], can be used, as implemented in BioJupies and iDep.

Table 2: Gene expression sources

| Application | User-provided data | GEO microarray | GEO RNA-seq |
|---|---|---|---|
| Gene pattern[2006] | + | + | - |
| GEO2R[2012] | - | + | - |
| BicOverlapper2 [2014] | + | + | - |
| Babelomics [2015] | + | - | - |
| Morpheus [2016] | + | - | - |

| Application | User-provided data | GEO microarray | GEO RNA-seq |
|---|---|---|---|
| START [2016] | + | - | - |
| BioJupies [2018] | + | - | + / - [1] |
| iDEP [2018] | + | - | + / - [1] |
| Degust [2019] | + | - | - |
| EXPANDER [2019] | + | - | - |
| GREIN [2019] | - | + | + |
| RaNa-seq [2019] | + | - | + |
| RNAdetector [2021] | + | - | - |
| GEOexplorer [2022] | - | + | - |
| **Phantasus** | + | + | + / - [1] |

Finally, we consider features that influence on user experience (Table 3). There are three major archetecture types used in gene expression analysis applications, which have different trade-offs between reactivity, ease of user set-up and ease of development: Shiny-based web-applications, non-Shiny web-applications and stand-alone appications. Some of non-Shiny web-applications (Gene pattern, BioJupies, RaNa-seq) allow to share user sessions, which insreases reproducibility and can be useful for collaborative working or to be used in publications. Interactive gene expression heatmaps and other plots help to visualize and explore the data, and are available in many tools. However the support of working with the metadata is limited. Only four other tools (Babelomic, Morpheus, EXPANDER, RNAdetector) allow to edit genes and sample annotations, and only Babelomic and iDEP can map gene annotations using external gene annotation databases.

[1] As processed in ARCHs4 and/or Dee2 projects

Table 3: User experience features

| Application | Architecture | Saved sessions | Interactive heatmap | Interactive plots | Editing sample annotations | Editing gene annotations |
|---|---|---|---|---|---|---|
| Gene pattern [2006] | web non-Shiny | + | - | - | - | - |
| GEO2R [2012] | web non-Shiny | - | - | - | - | - |
| BicOverlapper2 [2014] | local instalation | - | + / - [2] | - | - | - |
| Babelomics [2015] | web non-Shiny | - | - | + | + | + |
| Morpheus [2016] | web non-Shiny | + / - | + | + | + | - |
| START [2016] | web Shiny | - | + / - [2] | + | - | - |
| BioJupies [2018] | web non-Shiny | + | + / - [2] | + | - | - |
| iDEP [2018] | web Shiny | - | + / - [2] | + | - | + |
| Degust [2019] | web non-shiny | - | + | + | - | - |
| GREIN [2019] | web Shiny | - | + | + | - | - |
| EXPANDER [2019] | local instalation | - | + | + | + | - |
| RaNa-seq [2019] | web non-Shiny | + | + / - [2] | + | - | - |
| RNAdetector [2021] | local instaltation | - | - | - | + | - |
| GEOexplorer [2022] | web shiny | - | + | + | - | - |
| **Phantasus** | web non-Shiny | + | + | + | + | + |

Phantasus is a shiny-independent web-application that is able to perform all key steps of gene expression analysis and provide access to a huge collection of publicly available datasets, both microarray and RNA-seq. Using an interactive heatmap-based interface, users can explore the data, modify annotations, build plots etc. Moreover Phantasus provides results in publication-ready form: with image export to vector-based graphics formats and with session links, which keep all user-defined settings, such as colors, annotations and filter options.

[2] Limited number of genes
