## Supplementary File 2 for "Phantasus: web-application for visual and interactive gene expression analysis"

### **Supplementary File 2.** Case study: basic usage, full walk-through

#### Introduction

In this section we show example usage of Phantasus for analysis of public gene expression data from GEO database. It starts from loading data, normalization and filtering outliers, to doing differential gene expression analysis and downstream analysis.

To illustrate the usage of Phantasus let us consider public dataset from Gene Expression Omnibus (GEO) database GSE53986. This dataset contains data from experiments, where bone marrow derived macrophages were treated with three stimuli: LPS, IFNg and combined LPS+INFg.

#### Opening Phantasus

The simplest way to try Phantasus application is to go to web-site https://ctlab.itmo.ru/phantasus or its mirror https://artyomovlab.wustl.edu/phantasus where the latest versions are deployed. Alternatively, Phantaus can be installed locally (see section Stand-alone Phantasus distribution of the main arcticle).

When Phantasus opens the starting screen (Figure 1) should appear.

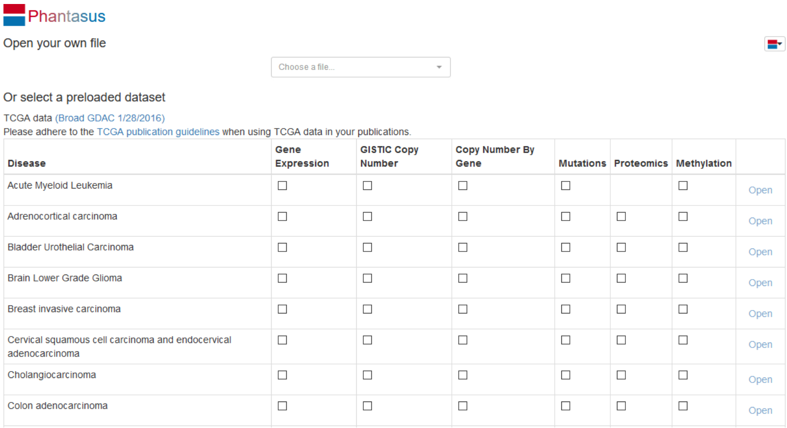


Figure 1: Phantasus starting screen

#### Preparing the dataset for analysis

##### Opening the dataset

Let us open the dataset. To do this, select *GEO Datasets* option in *Choose a file...* dropdown menu. There, a text field will appear where `GSE53986` should be entered. Clicking the *Load* button (or pressing *Enter* on the keyboard) will start the loading. After a few seconds, the corresponding heatmap (Figure 2) should appear.

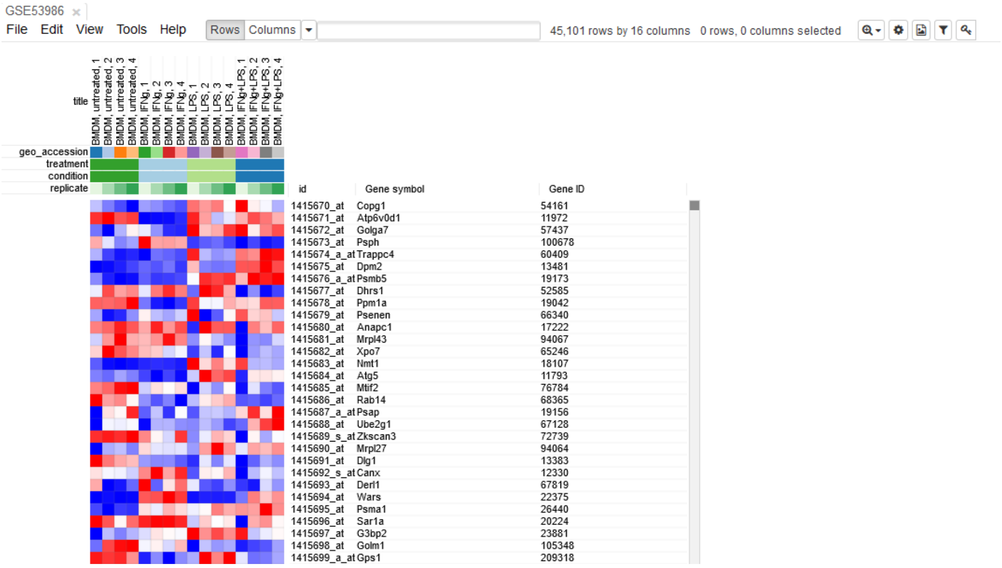


Figure 2: Heatmap of the loaded dataset

On the heatmap, the rows correspond to genes (or microarray probes). The rows are annotated with *Gene symbol* and *Gene ID* annotaions (as loaded from GEO database). Columns correspond to samples. They are annotated with titles, GEO sample accession identifiers and treatment field. The annotations, such as treatment, are loaded from user-submitted GEO annotations (they can be seen, for example, in *Charateristics* section at https://www.ncbi.nlm.nih.gov/geo/query/acc.cgi?acc=GSM1304836). We note that not for all of the datasets in GEO such proper annotations are supplied.

##### Adjusting expression values

By hovering at heatmap cell, gene expression values can be viewed. The large values there (Figure 3) indicate that the data is not log-scaled, which is important for most types of gene expression analysis.

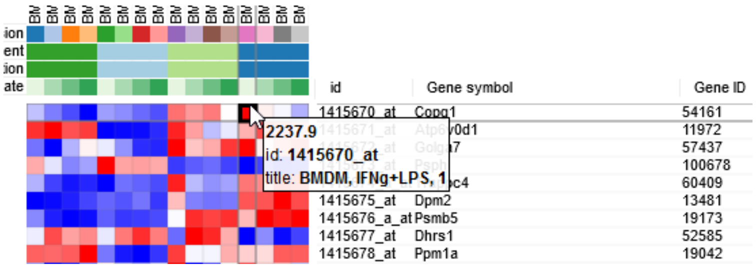


Figure 3: Heatmap cell tooltip

For the proper further analysis it is recommended to normalize the matrix. To normalize values go to *Tools/Adjust* menu and check *Log 2* and *Quantile normalize* adjustments (Figure 4).

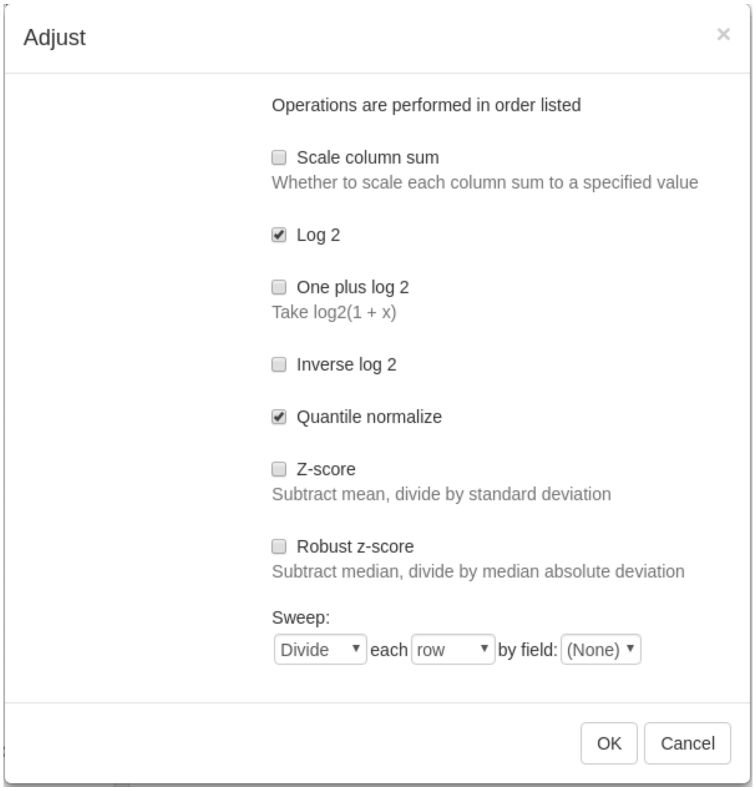


Figure 4: Adjust tool

The new tab with adjusted values will appear. All operations that modify gene expression matrix (such as adjustment, subsetting and several others) create a new tab. This allows to revert the operation by going back to one of the previous tabs.

#### Removing duplicate genes

Since the dataset is obtained with a mircroarray, a single gene can be represented by several probes. This can be seen, for example, by sorting rows by *Gene symbol* column (one click on column header), entering `Actb` in the search field and going to the first match by clicking down-arrow next to the field. There are five probes corresponding to Actb gene in the considered microarray (Figure 5).

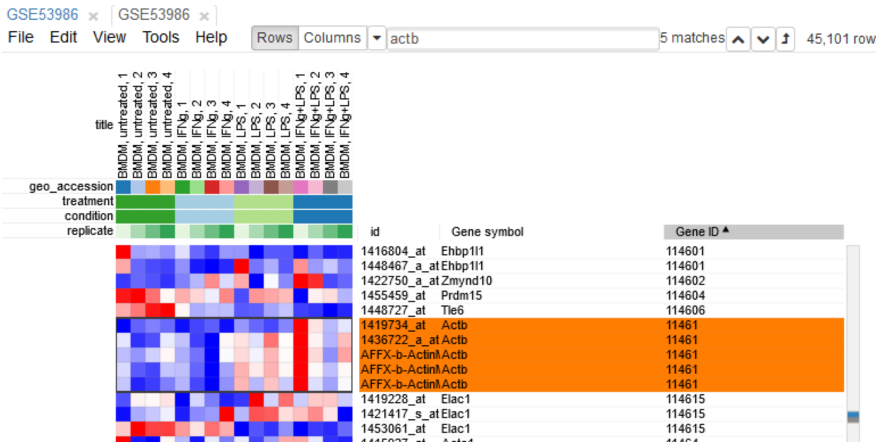


Figure 5: `Actb` duplicates

To simplify the analysis it is better to have one row per gene in the gene expression matrix. One of the easiest ways is to chose only one row that has the maximal median level of expression across all samples. Such method removes the noise of lowly-expressed probes. Go to *Tools/Collapse* and choose *Maximum Median Probe* as the method and *Gene ID* as the collapse field (Figure 6).

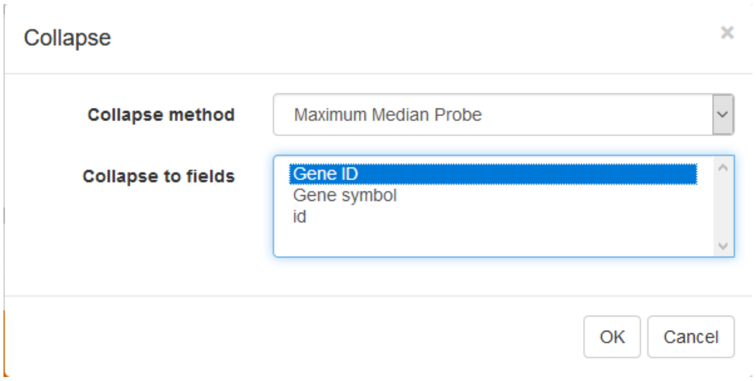


Figure 6: Collapse tool settings

The result will be shown in a new tab.

#### Filtering lowly-expressed genes

Additionally, lowly-epxressed genes can be filtered explicitly. It helps to reduce noise and increase power of downstream analysis methods.

First, we calculate mean expression of each gene using *Tools/Create Calculated Annotation* menu. Select `Mean` operation, optionally enter a name for the resulting column (Figure 7), and click *OK*. The result will appear as an additional column in row annotations (Figure 8).

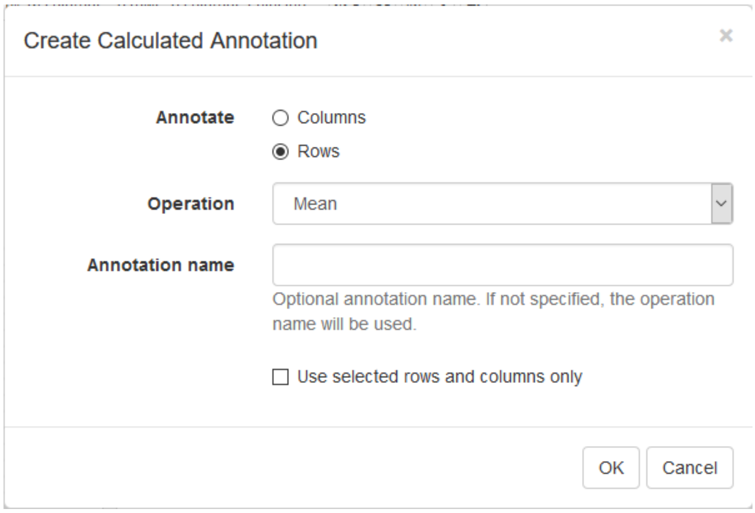


Figure 7: Row mean calculated annotation settings

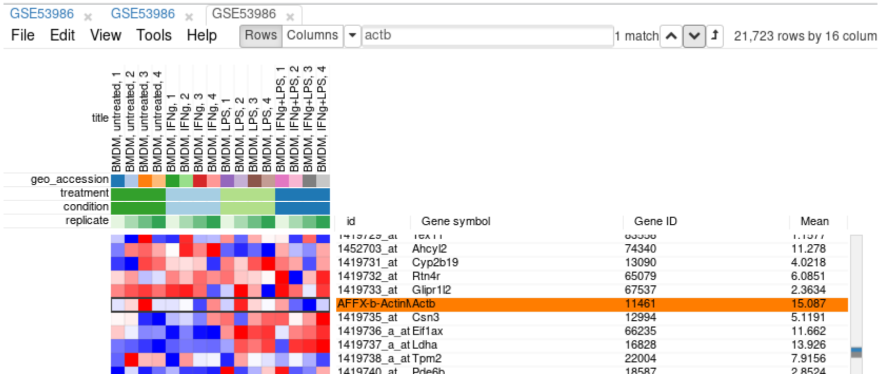


Figure 8: Heatmap with loaded row mean annotation

Now this annotation can be used to filter genes. Open *Tools/Filter* menu. Click *Add* to add a new filter. Choose `mean_expression` as a *Field* for filtering. Then press *Switch to top filter* button and input the number of genes to keep. A good choice for a typical mammalian dataset is to keep around 10–12 thousand most expressed genes (Figure 9). Filter is applied automatically, so after closing the dialog with *Close* button only the genes passing the filter will be displayed (Figure 10).

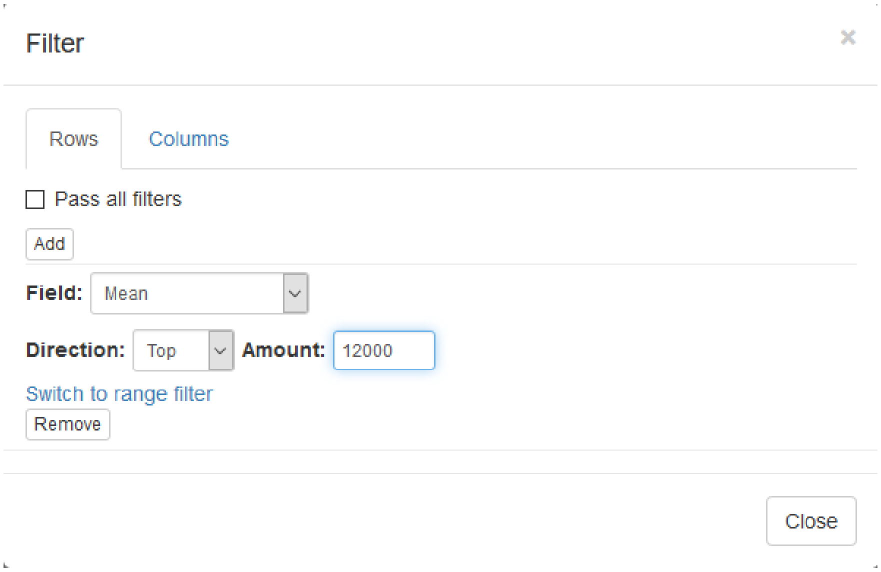


Figure 9: Filter tool settings

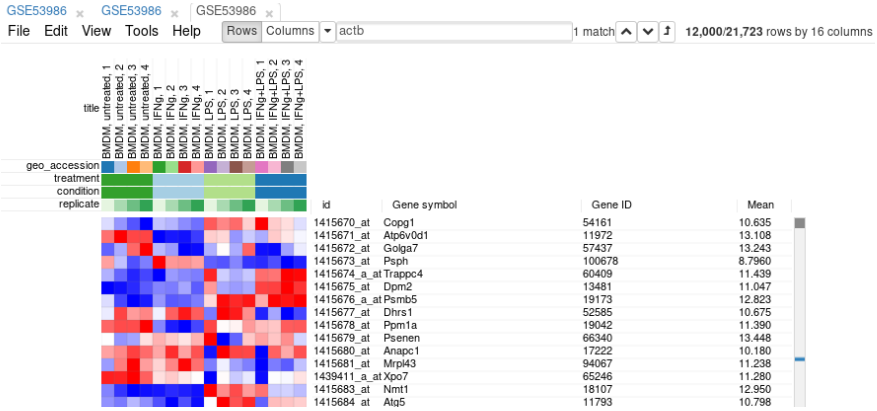


Figure 10: Heatmap with filtered genes

It is more convenient to extract these genes into a new tab. For this, select all genes (click on any gene and press *Ctrl+A*) and use *Tools/New Heat Map* menu (or press *Ctrl+X*).

Now you have the tab with a fully prepared dataset for the further analysis. To easily distinguish it from other tabs, you can rename it by right click on the tab and choosing *Rename* option. Let us rename it to `GSE53986_norm`.

It is also useful to save the current result to be able to return to it later. In order to save it use

*File/Save Dataset* menu. Enter an appropriate file name (e.g. `GSE53986_norm`) and press `OK`. A file in text GCT format will be downloaded.

### Exploring the dataset

#### PCA Plot

One of the ways to asses quality of the dataset is to use principal component analysis (PCA) method. This can be done using *Tools/Plots/PCA Plot* menu (Figure 11).

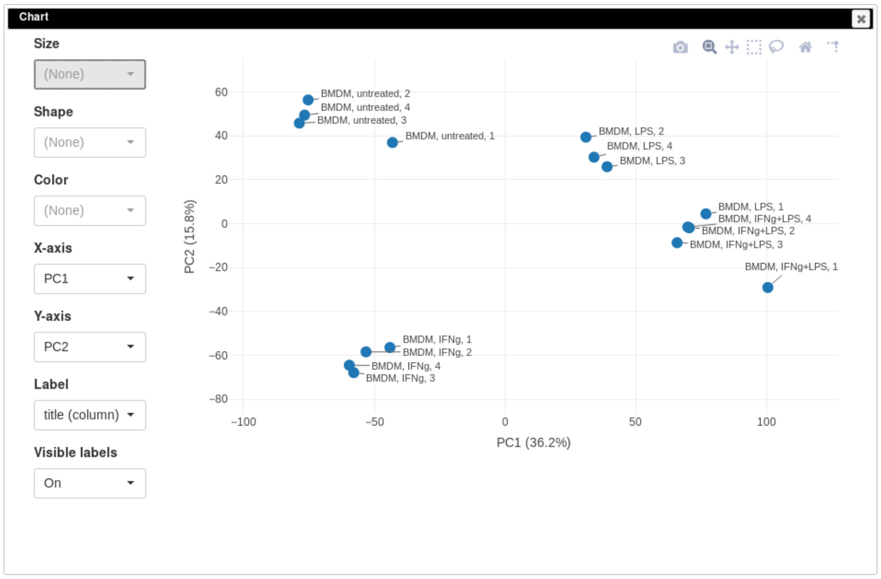


Figure 11: PCA plot for dataset GSE53986

You can customize color, size and labels of points on the chart using values from annotation (Figure 12). Here we set color to come from *treatment* annotation.

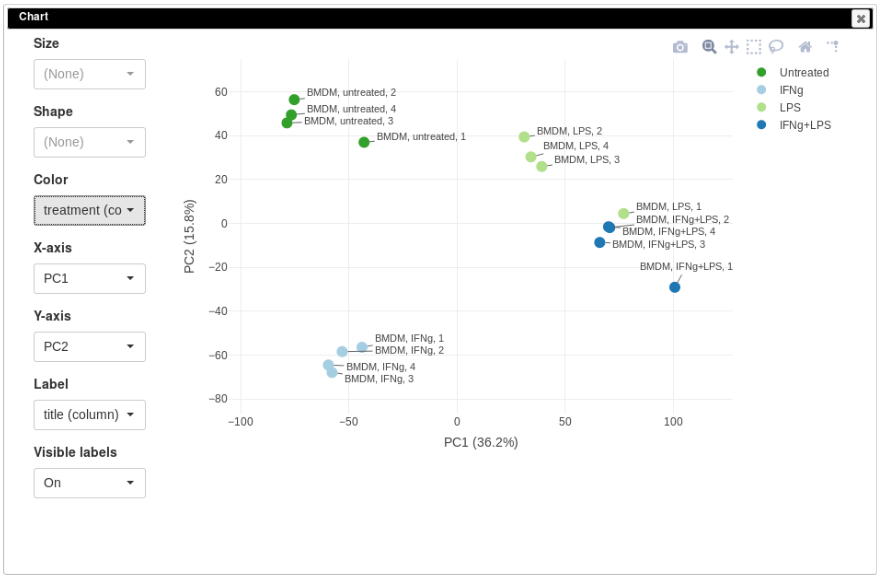


Figure 12: Custimized PCA plot for the dataset GSE56986

It can be seen that in this dataset the first replicates in each condition are outliers.

#### K-means clustering

Another useful dataset exploration tool is k-means clustering. Use *Tools/Clustering/k-means* to cluster genes into 16 clusters (Figure 13).

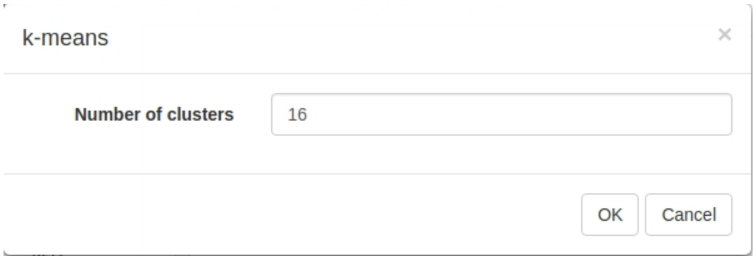


Figure 13: K-means tool settings

Afterwards, rows can be sorted by *clusters* column. By using menu *View/Fit to window* one can get a "bird's-eye view" on the dataset (Figure 14). Here also one can clearly see outlying samples.

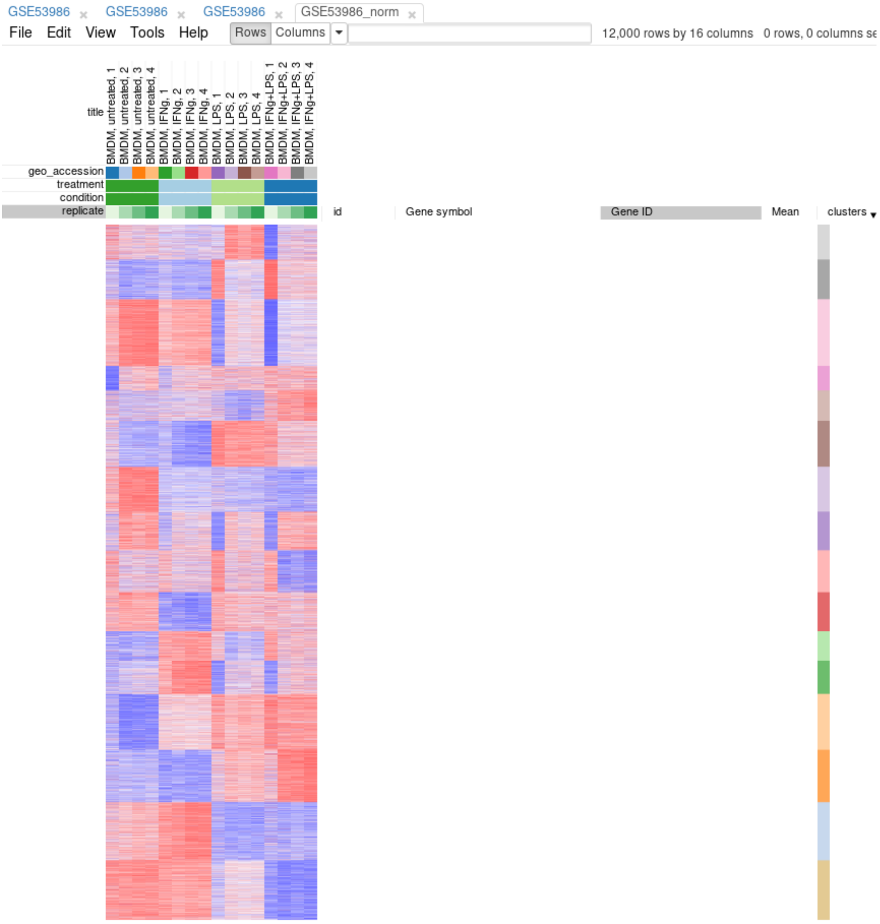


Figure 14: Clusters obrained using the K-means algorithm

#### Hierarchical clustering

*Tool/Hierarchical clustering* menu can be used to cluster samples and highlight outliers (and concordance of other samples) even further (Figure 15).

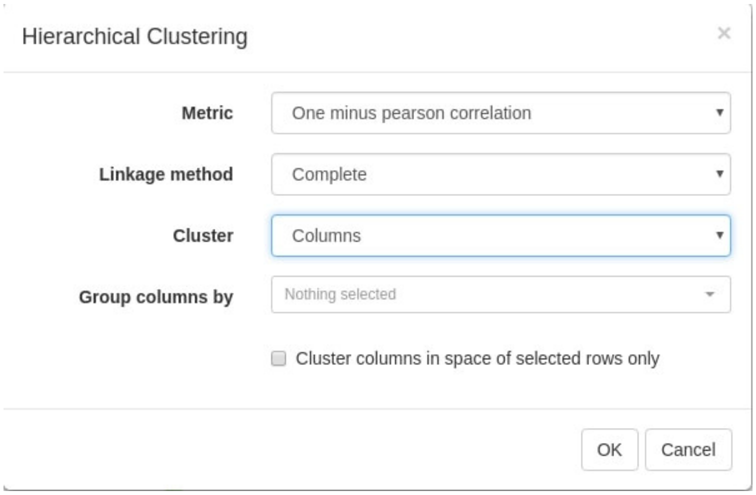


Figure 15: Hierarchical clustering settings

#### Filtering outliers

Now, when outliers are confirmed and easily viewed with the dendrogram from the previous step, you can select the good samples (Figure 16) and extract them into another heatmap (by clicking *Tools/New Heat Map* or pressing *Ctrl+X*).

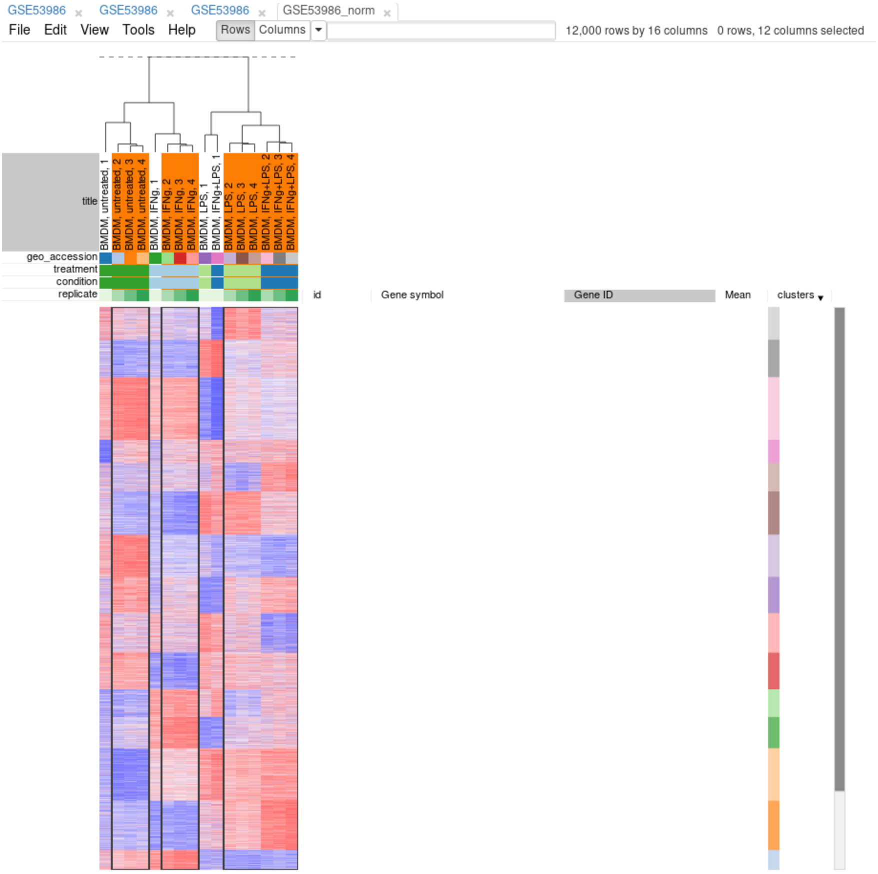


Figure 16: Good samples selection

#### Differential gene expression

##### Apllying *limma* tool

Differential gene expression analysis can be carried out with *Tool/Diffential Expression/limma* menu. Choose *treatment* as a *Field*, with *Untreated* and *LPS* as classes (Figure 17). Clicking *OK* will call differential gene expression analysis method with *limma* R package.

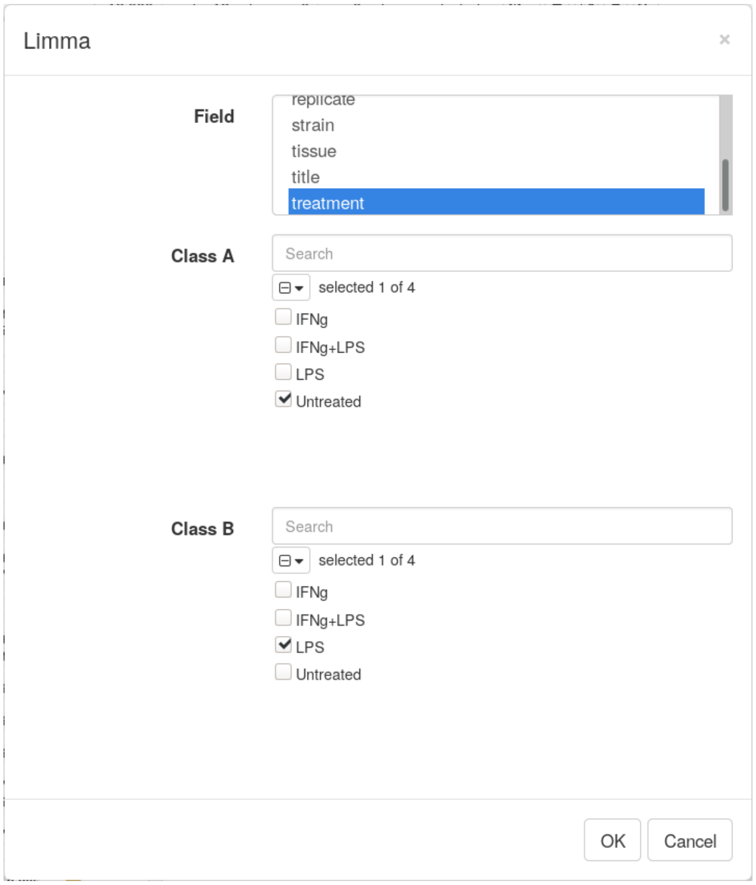


Figure 17: Settings for differential expression analysis by `limma` package

The rows can be ordered by decreasing *t*-statistic column to see which genes are the most up-regulated upon LPS treatment (Figure 18).

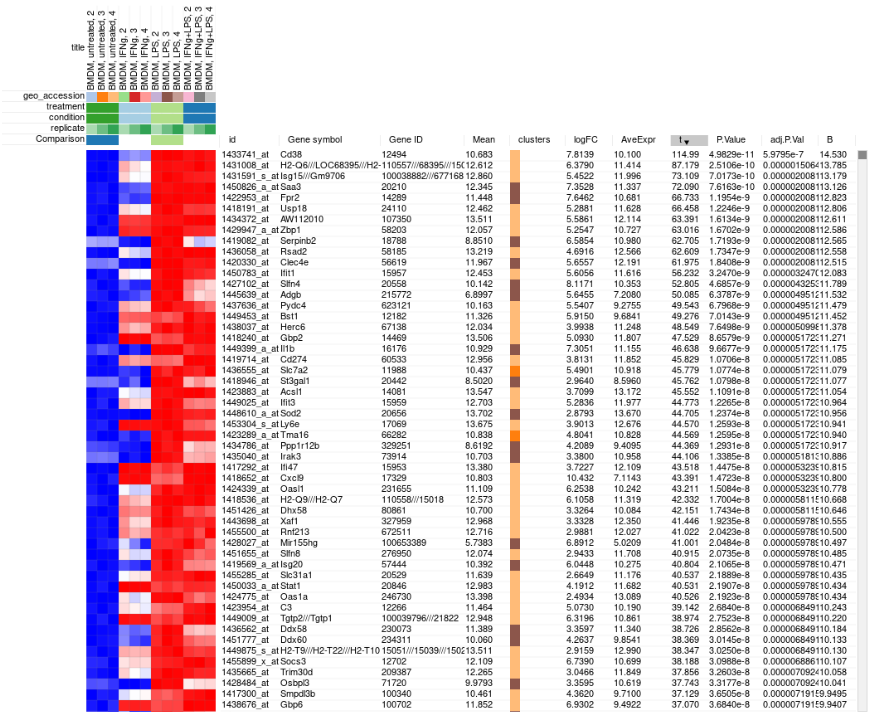


Figure 18: Differential expression analysis results for the dataset GSE53986

#### Pathway analysis with FGSEA

The results of differential gene expression can be used for pathway enrichment analysis with *FGSEA* tool.

Open *Tools/Pathway Analysis/Perform FGSEA*, then select Pathway database, which corresponds specimen used in dataset (Mus Musculus in this example), ranking column and column with ENTREZID or Gene IDs (Figure 19).

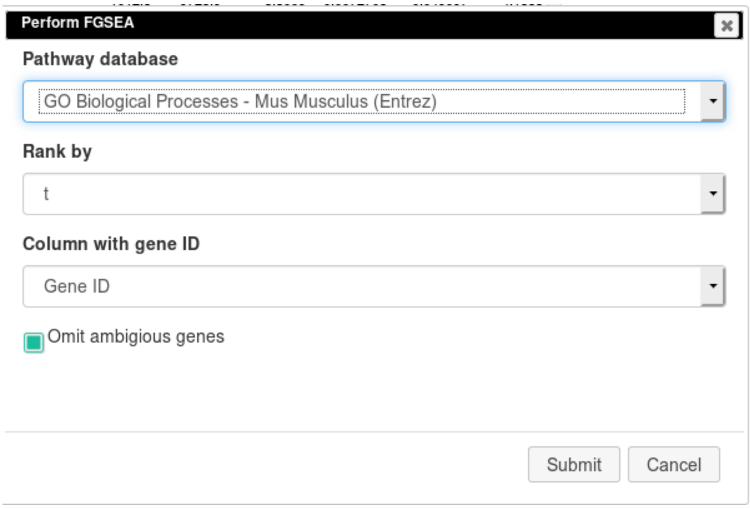


Figure 19: FGSEA settings

Clicking OK will open new tab with pathways table (Figure 20).

**Actions:**

Save as TSV

**FGSEA:**

| pathway | pval | padj | log2err | ES | NES | size | leadingEdge |
|---|---|---|---|---|---|---|---|
| Defense response to virus | 1.00e-10 | 4.06e-8 | NA | 0.681 | 2.67 | 152 | 56417 15958 58185 20846 80861 ... |
| Immune system process | 1.00e-10 | 4.06e-8 | NA | 0.523 | 2.29 | 398 | 56619 16452 16365 213233 56417 ... |
| Innate immune response | 1.00e-10 | 4.06e-8 | NA | 0.546 | 2.33 | 313 | 56619 16452 16365 56417 15958 ... |
| Response to bacterium | 1.00e-10 | 4.06e-8 | NA | 0.573 | 2.28 | 175 | 18126 16365 20846 20558 80861 ... |
| Immune response | 1.79e-10 | 5.83e-8 | 0.827 | 0.525 | 2.17 | 213 | 21938 56619 17130 20304 23962 ... |

Figure 20: Pathways enriched by FGSE tool for the dataset GSE54986

Clicking on table row will provide additional information on pathway: pathway name, genes in pathway, leading edge. You can save result of analysis in `TSV` format.
