## Supplementary File 3 for "Phantasus: web-application for visual and interactive gene expression analysis"

### **Supplementary File 3.** Case study: data reanalysis, full walk-through

#### Overview

This section covers the replication of the stuby by Mowel and colleagues [Mowel et al., 2017]. In the study the authors show that Rroid locus control innate lymphoid cells (ILCs) by promoting *Id2* gene expression. Here we consider two stages of the analysis which can be performed using Phantasus:

- obtaining *Id2*-dependent gene signature based on the dataset GSE76466;
- establishing *Rroid*-mediated regulation of the signature in the dataset GSE101459.

During the section we will provide screenshots to demonstrate controls settings for replication.

#### Obtaining Id2-dependent gene signature

In the original paper the authors use GSE76466 [Shih et al., 2016] dataset that contains wild-type (WT) and *Id2*-null natural killer (NK) cells samples to determine *Id2*-dependent signature. Using Phantasus this dataset can be explored in a straightforward manner.

As a first step of the analysis we will explore dataset. The dataset can be opened directly in Phantasus by its GEO identifier. After the dataset is loaded we can make a principal component analysis (PCA) plot to get an overview of the dataset. For the analysis we will keep only WT and *Id2*-null samples and the top 12000 expressed genes. Furthermore we perform lograithmic and quantile normalization to scale the raw counts.

The obtained PCA plot Figure 1 shows clear separation between WT and *Id2*-null samples. This encourage us to do next step and determine genes that are differentially expressed between this two genotypes.

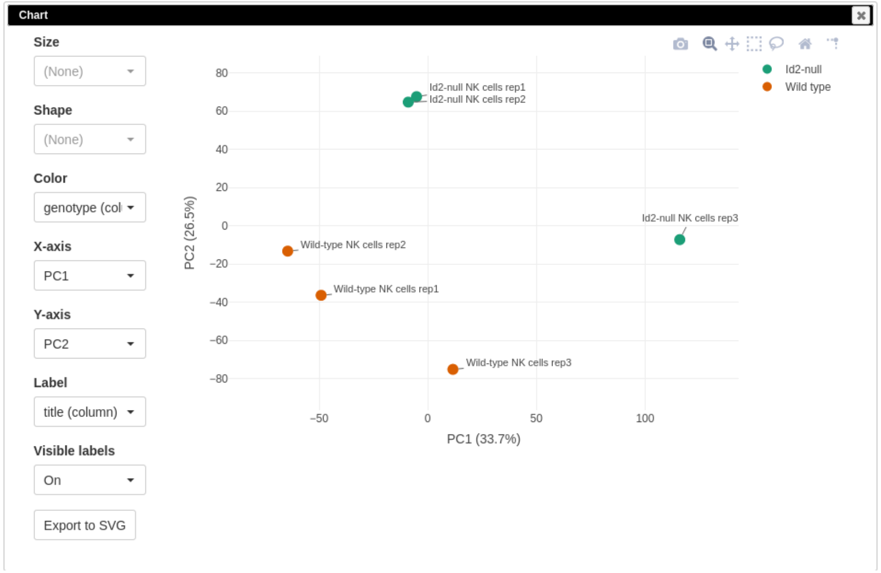


Figure 1: PCA plot for dataset GSE76466

Following the authors we will use DESeq2 tool [Love et al., 2014] for differential gene expression analysis. Notice that DESeq2 requires raw count matrix with all the genes as an input, so all of the further steps are performed with the original dataset, but only for the WT and *Id2*-null samples. A volcano plot done with *Plots/Volcano Plot* function gives an overview of differential expression Figure 2. Notably, *Id2* mRNA expression is down-regulated in the knockout samples, which serves as a positive control.

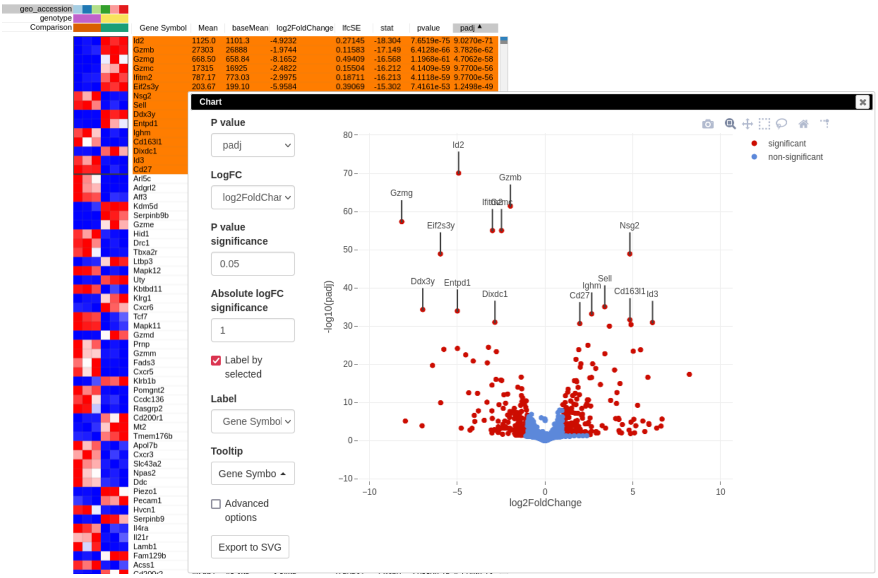


Figure 2: Volcano plot with differential gene expression results between WT and *Id2*-deficient samples for dataset GSE76466

Desirable *Id2*-dependent gene signature, used by the authors, consists of the genes up-regulated in *Id2*-null samples compared to WT. To obtain this siguntature we can apply `padj < 0.05` and `stat >= 0` filters using *Tools/Filter* menu. Then we copy gene symbols from the filtered rows and paste them into a text file for the next stage.

#### ***Rroid*-mediated regulation in group 1 ILCs**

Mowel with colleauges suggested that the discovered *Rroid* locus regulation of ILCs development happens via regulation of *Id2* gene. In order to computationally confirm this hypothesis, the authors examined the enrichment of the previously obtained gene signature in the dataset with *Rroid*-deficient and wild-type samples. The *Rroid* dataset generated by the authors is available at GEO database under accession number GSE101459.

After dataset GSE101459 is loaded in Phantasus we first can make a PCA plot, by applying the same steps as we did for the dataset GSE76466: filtration of lowly expressed genes, logarithmic and quantile normalization. The PCA plot Figure 3 indicates that the first samples in each of the two groups differ signficantly from the rest, which could indicate that they are outliers. However the samples are separated by the genotype, so we can use the whole dataset for the further analysis.

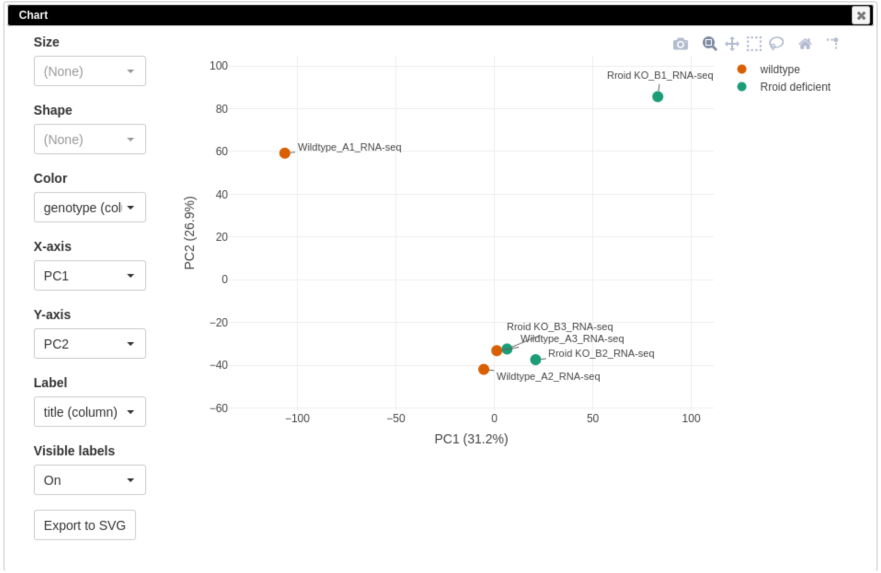


Figure 3: PCA plot for dataset GSE101459

Again, as we consider an RNA-seq dataset, the differential expression analysis (DE) is performed using DESeq2 on the original count values. Resulting volcano plot Figure 4 reveals significant down-regulation of Id2 gene, which is consistent with the main hypothesis.

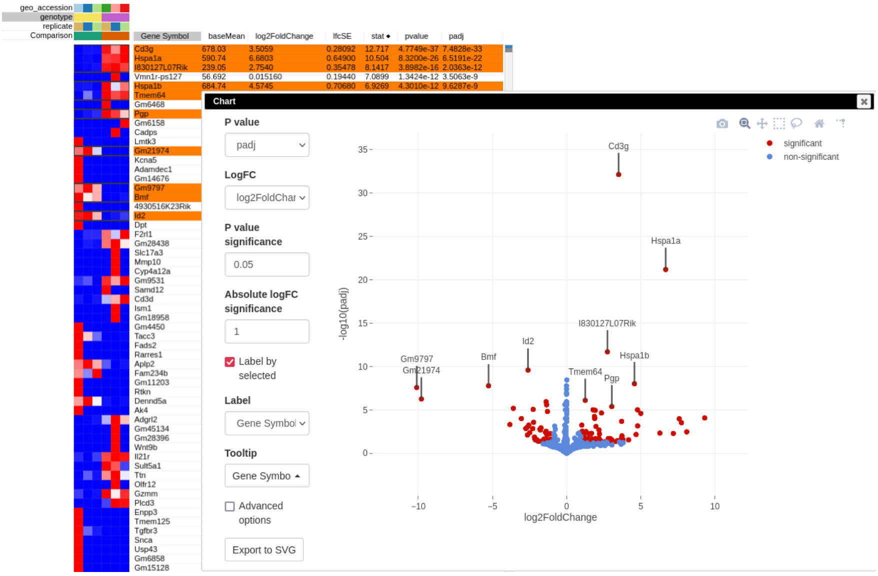


Figure 4: Volcano plot with differential gene expression results between WT and *Rroid*-deficient samples for dataset GSE101459

Following Mowel et al [Mowel et al., 2017] we want to perform gene set enrichment analysis (GSEA). GSEA requires to rank genes by the differense in the expression. For GSEA analysis we are going to use only the top 12000 expressed genes (filtered by `baseMean` column). In addition we scale gene expression values using logarithmic and quatile normalization, so that we have a cleaner heatmap representation of the gene expression matrix.

Finally, to make a GSEA plot Figure 5 we copy the saved gene signature obtained earlier Section , paste it into search field of the filtered DE results, and use *Tools/Plots/GSEA Plot* menu. The resulting plot shows that the *Id2*-dependent genes are up-regulated in the *Rroid*-deficient samples. This plot replicates figure 4E of the original paper by Mowel and colleagues.

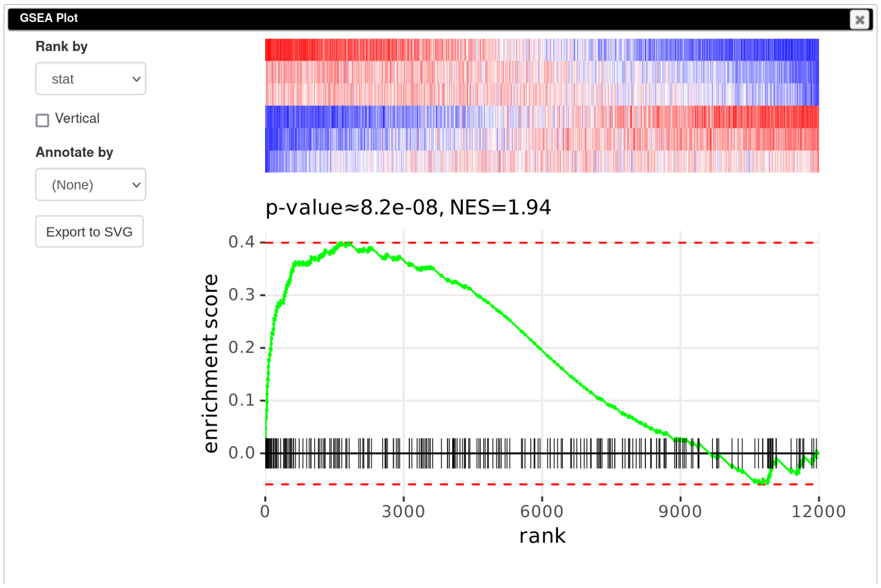


Figure 5: Enrichment plot of *Id2*-dependent gene signature in dataset GSE101459

### References

Walter K. Mowel, Sam J. McCright, Jonathan J. Kotzin, Magalie A. Collet, Asli Uyar, Xin Chen, Alexandra DeLaney, Sean P. Spencer, Anthony T. Virtue, EnJun Yang, Alejandro Villarino, Makoto Kurachi, Margaret C. Dunagin, Gretchen Harms Pritchard, Judith Stein, Cynthia Hughes, Diogo Fonseca-Pereira, Henrique Veiga-Fernandes, Arjun Raj, Taku Kambayashi, Igor E. Brodsky, John J. O'Shea, E. John Wherry, Loyal A. Goff, John L. Rinn, Adam Williams, Richard A. Flavell, and Jorge Henao-Mejia. Group 1 innate lymphoid cell lineage identity is determined by a cis-regulatory element marked by a long non-coding rna. *Immunity*, 47(3):

435–449.e8, 2017. ISSN 1074-7613. doi: https://doi.org/10.1016/j.immuni.2017.08.012. URL https://www.sciencedirect.com/science/article/pii/S1074761317303709.

Han-Yu Shih, Giuseppe Sciumè, Yohei Mikami, Liying Guo, Hong-Wei Sun, Stephen R. Brooks, Joseph F. Urban, Fred P. Davis, Yuka Kanno, and John J. O'Shea. Developmental acquisition of regulomes underlies innate lymphoid cell functionality. *Cell*, 165(5):1120–1133, 2016. ISSN 0092-8674. doi: https://doi.org/10.1016/j.cell.2016.04.029. URL https://www.sciencedirect.com/science/article/pii/S0092867416304238.

Michael I. Love, Wolfgang Huber, and Simon Anders. Moderated estimation of fold change and dispersion for rna-seq data with deseq2. *Genome Biology*, 15, 2014.
